# A Folding–Docking–Affinity framework for protein–ligand binding affinity prediction

**DOI:** 10.1101/2024.04.16.589805

**Authors:** Ming-Hsiu Wu, Ziqian Xie, Degui Zhi

## Abstract

Accurate protein-ligand binding affinity prediction is crucial in drug discovery. Existing methods are predominately docking-free, without explicitly considering atom-level interaction between proteins and ligands in scenarios where crystallized protein-ligand binding conformations are unavailable. Now, with breakthroughs in deep learning AI-based protein folding and binding conformation prediction, can we improve binding affinity prediction? This study introduces a framework, Folding-Docking-Affinity (FDA), which folds proteins, determines protein-ligand binding conformations, and predicts binding affinities from three-dimensional protein-ligand binding structures. Our experimental results indicate that FDA performs comparably to state-of-the-art docking-free methods. We anticipate that our proposed framework serves as a starting point for integrating binding structures for more accurate binding affinity prediction.

## 1 Introduction

Protein-ligand binding affinity prediction is an important problem in drug discovery and computational biology. Accurately predicting the binding affinity between a protein and a small molecule ligand can help identify potential drug candidates and optimize drug design. Most existing binding affinity prediction methods leverage machine-learning methods, but they do not typically consider the binding pose, i.e., docking-free, hence ignoring the atom-level interactions between ligands and proteins, as shown in Fig. 1(a). In these models, proteins are primarily represented as amino acid sequences or protein contact map graphs while SMILES strings or molecular graphs represent ligands. Deep neural networks are then applied to extract important latent features for predicting binding affinity (1; 2; 3; 4; 5; 6; 7; 8; 9). The classical models include DeepDTA (10), GraphDTA (11), DGraphDTA (12), and MGraphDTA (13). Additionally, for the sub-problem of kinase-drug binding affinity prediction, a specialized model, KDBNet, has also been proposed (14). It differs from the aforementioned models in that it incorporates features from predefined three-dimensional kinase binding pocket, thereby implicitly encodes the binding pose information. But this approach is specifically designed for the kinase-ligand binding affinity and may not be generalizable to wider-ranges of proteins. These docking-free methods often function as black-box models lacking structural context. Although they can be easily trained in an end-to-end manner, these docking-free methods may lack the interpretability and detailed insight into molecular interactions provided by docking approaches, potentially leading to less informed predictions.

**Figure 1:**
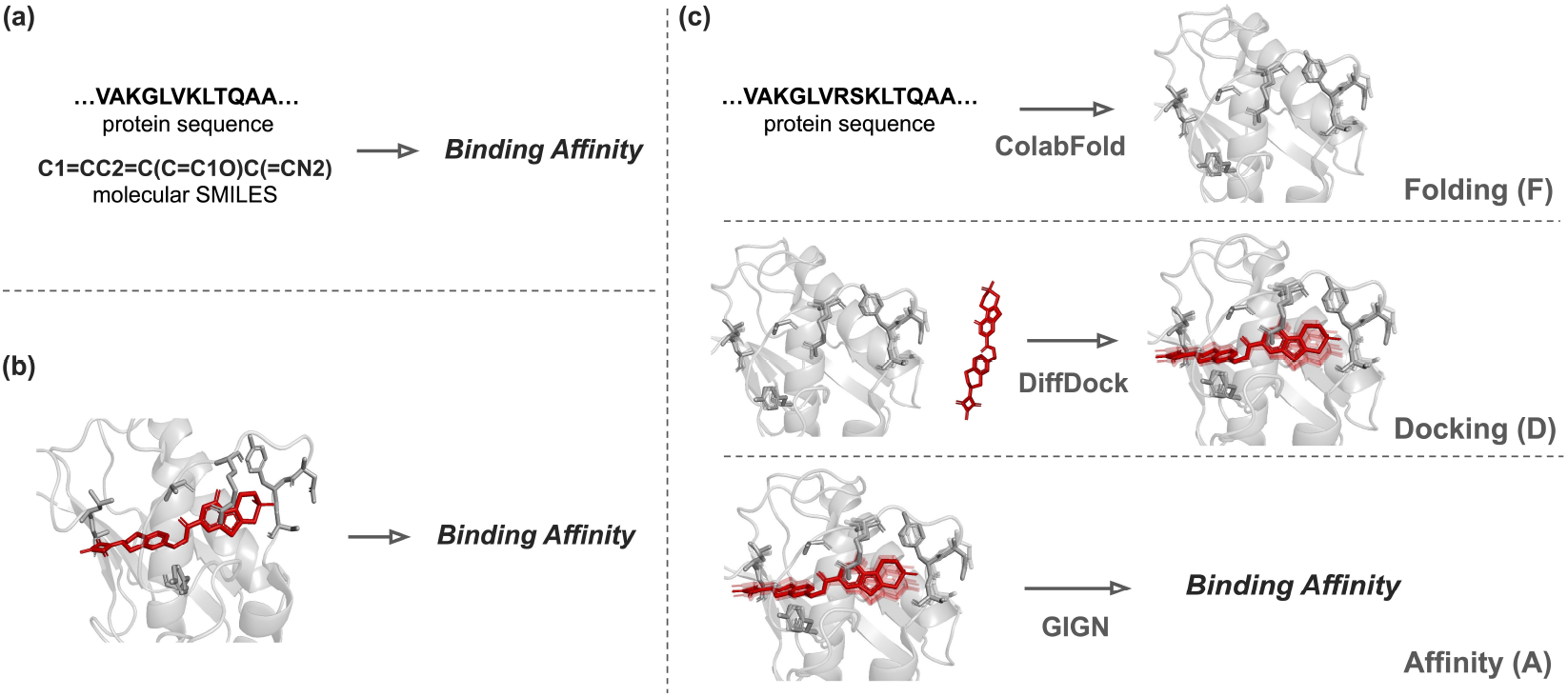
Comparison of Docking-Free, Docking-Based, and FDA Methods for Binding Affinity Prediction. (a) Docking-free models predict binding affinity directly from a protein sequence and a molecular SMILES. (b) Docking-based models predict binding affinity directly from the experimentally determined crystallized binding structures. (c) Our work, Folding-Docking-Affinity (FDA), consists of three components: folding proteins (Folding), determining optimal protein-ligand binding poses (Docking), and predicting binding affinities from computed three-dimensional protein-ligand binding structures (Affinity). The crystal structure of the protein-ligand complex (PDB ID: 3DPF) (56; 44) is used for illustration.

Meanwhile, when co-crystallized 3D structures are available, there has been notable progress in the development of docking-based approaches, shown in Fig. 1(b), that consider atom-level interactions explicitly (15; 16; 17; 18; 19; 20; 21; 22), exemplified by works like SchNet (23), EGNN (24), and GIGN (25). These works utilize 3D convolutional neural networks or interaction graph neural networks to predict binding affinity. Notably, these docking-based models could outperform their docking-free counterparts when testing on the PDBBind dataset (26) where a collection of high-resolution crystallized three-dimensional binding structures are curated. It is crucial to highlight, however, that this superiority would depend upon the availability of the three-dimensional binding structures. A pertinent question arises: in the absence of a high-resolution binding structure, can the docking-based model still maintain its advantage? This work aims to answer this question.

The premises of our question are the recent breakthroughs in deep learning-based protein folding and docking. Given AlphaFold (27; 28) and other methods (29; 30; 31; 32; 33; 34; 35) have revolutionized the field of protein folding, accessing three-dimensional protein structures become much easier than before. Simultaneously, deep learning-based protein-ligand docking models (36; 37; 38; 39) have been developed to generate binding conformations when provided with specific protein and ligand information much more easily, and their accuracy is catching up with computationally intensive molecular dynamics based docking. These advancements pave the way for a new approach to predict binding affinity in situations where a high-resolution crystal three-dimensional binding structure is not available.

Here, we present our framework, Folding-Docking-Affinity (FDA), illustrated in Fig. 1(c), designed to fold proteins, determine optimal protein-ligand binding poses, and then predict binding affinity from computed three-dimensional binding structures. Notably, our approach is versatile and applicable to any protein-ligand pair. Here, we utilized ColabFold (31), a protein structure prediction model, DiffDock (36), a state-of-the-art deep learning-based docking model, and a GNN-based affinity predictor, GIGN (25), to build this framework. Importantly, any component of this framework can be substituted with alternative models. We expect that this approach enables reliable predictions of binding affinities of protein-ligand pairs without crystallized binding structures. Our assumption is that the docking-based method could outperform its docking-free counterparts by considering atom-level interactions, which more accurately reflect the true physical dynamics, even with a minor deviation from the ground truth.

Our contributions are: We construct a general protein-ligand affinity prediction framework based on a computed explicit binding pose. We design the framework with individual replaceable components and thus can adapt to the fast-moving pace of folding, docking, and affinity prediction methods development. The FDA exhibits performance comparable to SOTA docking-free methods in kinase-specific benchmark experiments using the DAVIS (40) and KIBA (41) datasets, demonstrating its feasibility for binding affinity prediction. Additionally, we observed that accidental noise introduced during the folding and docking steps can unexpectedly enhance binding affinity prediction performance. Inspired by this, we incorporated generated binding poses from these stages as a data augmentation strategy, further improving predictive performance.

## 2 Results

### 2.1 Framework

This study aims to develop a method for predicting binding affinity by explicitly considering proteinligand binding conformations, when high-resolution co-crystallized structures are unavailable. We propose an end-to-end framework, FDA, which bridges the protein amino acid sequence and ligand to its corresponding binding affinity with computed binding conformations. The framework consists of three key components, as shown in Fig. 1(c). (i) Folding: Generating three-dimensional protein structures from protein amino acid sequences. (ii) Docking: Docking the ligand onto the generated protein structure. (iii) Affinity: Predicting binding affinities from predicted three-dimensional proteinligand binding structure. Importantly, our framework features individual replaceable components, allowing it to adapt to various folding, docking, and affinity prediction methods.

### 2.2 Affinity prediction benchmark

Our FDA method was benchmarked on two distinct public kinase-specific datasets, DAVIS (40) and KIBA (41), to evaluate its binding affinity prediction performance. Additionally, to assess its generalizability, both datasets were divided into four different test scenarios: both-new, new-drug, new-protein, and sequence-identity split. The results, presented in Fig. 2, are evaluated using the Pearson correlation coefficient (*R*_*p*_) and Mean Squared Error (MSE) metrics. The detailed numerical data can be found in Table. S1-S2. In the both-new split, our FDA method outperforms its docking-free counterparts in most cases, showing *R*_*p*_ values of 0.29 and 0.51 in the DAVIS and KIBA datasets, respectively. However, in the MSE metric for the DAVIS dataset, DGraphDTA exhibits the lowest MSE. In the new-drug split, the performance of FDA is on par with MGraphDTA in the DAVIS dataset, with *R*_*p*_ values of 0.34. For the KIBA dataset, the FDA method slightly outperforms MGraphDTA and DGraphDTA. In the new-protein and sequence-identity splits, the FDA method surpasses other docking-free models in the DAVIS dataset. However, in the KIBA dataset, MGraphDTA achieves the best performance in both *R*_*p*_ and MSE, while our FDA method performs comparably to DGraphDTA.

**Figure 2:**
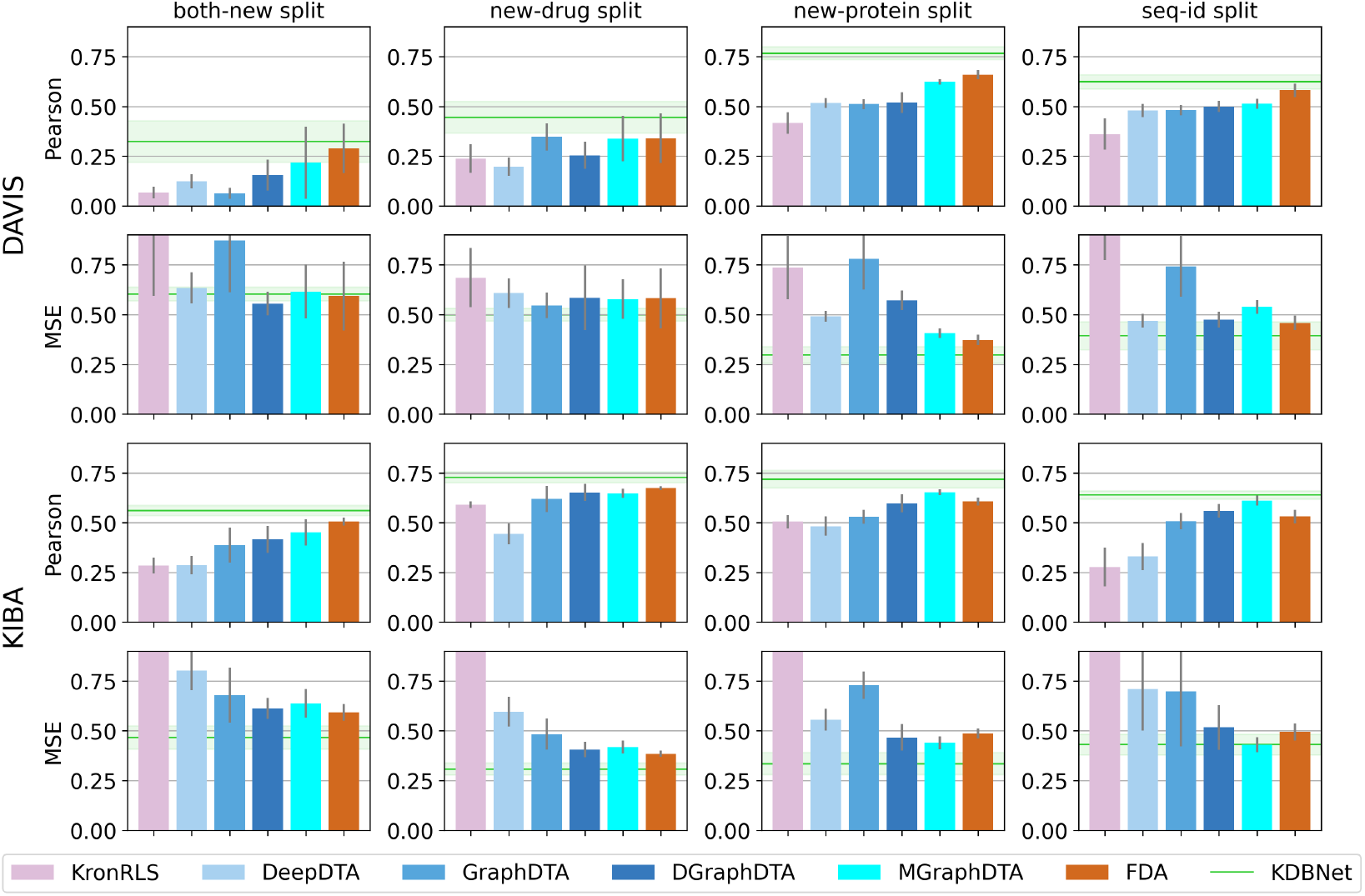
Performance Comparison of FDA and Docking-Free Methods on DAVIS and KIBA Datasets. Bar plots display the average *±* standard deviation of the evaluation results across five randomly chosen train/test splits. The light green horizontal line and area represent the average and the standard deviation of KDBNet. Pearson correlation coefficient (*R*_*p*_) and Mean Squared Error (MSE) were calculated based on the predicted and true *pK*_*d*_ values. The experimental results of KronRLS, DeepDTA, GraphDTA, DGraphDTA, and KDBNet are extracted from (14). The detailed numerical data can be found in Table. S1-S2.

We observed that MGraphDTA performs well on the random split, as illustrated in Fig. S1, where proteins and ligands in the test set have some overlap with the training set, its performance, however, significantly declines in the more challenging scenarios. This drop in the performance of the docking-free model indicates potential overfitting and low generalizability. The same trend was also observed in the prior research (25). These findings underscore the advantage of our approach, which explicitly accounts for the protein-ligand binding conformations, in enhancing model generalizability when compared to other docking-free methods. Additionally, we found that, within our FDA framework, the use of ColabFold-generated apo-structures, in comparison to crystallized holo-structures —yielded a surprising improvement in affinity prediction performance. This enhancement was mostly observed in both *R*_*p*_ and MSE metrics, with the exception of the both-new split, as illustrated in Fig. S2.

The performance of KDBNet is also presented in Fig. 2 for a reference comparison. This kinase-specific model, which integrates features from a predefined three-dimensional kinase binding pocket, surpasses our method and other docking-free models across the four distinct split test sets of the DAVIS and KIBA datasets. We argue that the superior performance of KDBNet can be attributed to the utilization of well-identified and extensively studied kinase binding pockets. In contrast, our method and other docking-free models operate without the constraints of a predefined binding pocket, which may introduce a disadvantage in this comparison. Nonetheless, the performance of KDBNet establishes a benchmark for these datasets, highlighting the potential advantages of incorporating detailed structural information into predictive models.

### 2.3 Ablation study

In our affinity prediction benchmark using the DAVIS and KIBA dataset, where we tested the FDA method alongside other docking-free methods, our FDA approach demonstrates performance on par with SOTA docking-free models. However, the impacts of the deviation introduced by ColabFold and DiffDock on the final affinity prediction remain unclear. To address this, we designed ablation experiments with three distinct scenarios: (i) Crystal binding poses (Crystal-Crystal): Protein-ligand binding structures are determined solely through experiments, with no deviation introduced by ColabFold and DiffDock. (ii) Holo Crystal Proteins with DiffDock (Crystal-DiffDock): In this scenario, we utilize holo Crystal Proteins for protein structures and employ DiffDock to determine the ligand binding pose. (iii) Apo ColabFold with DiffDock (ColabFold-DiffDock): In this case, we use apo ColabFold protein structures and again utilize DiffDock to find the ligand binding pose. Details of the experimental setting can be found in Methods 5.3. Given the limited availability of crystallized binding structures for most protein-ligand pairs in the DAVIS dataset, we utilized the PDBBind (version 2016) dataset (26) with the three settings mentioned above to train three distinct affinity prediction models: Crystal-Crystal, Crystal-DiffDock, and ColabFold-DiffDock. After training, we assessed the performances of these models using a test set (DAVIS-53) selected from the DAVIS dataset. This test set consists of protein-ligand pairs with crystallized binding structures so that we can similarly curate three test sets under our three settings.

The performance of three distinct models on various test sets is illustrated in Fig. 3 and Table. S3. Initially, we hypothesize that the model trained using the Crystal-Crystal dataset could exhibit the best prediction performance on the Crystal-Crystal test set (indicated by the lightest color), while the ColabFold-DiffDock model predicting the ColabFold-DiffDock dataset could show the worst performance (represented by the darkest color). This assumption is based on the potential error introduced by ColabFold and DiffDock, which may impact the model’s performance. On the contrary, our observations reveal that the lowest Root Mean Square Error (RMSE) is observed in the Crystal-Crystal and ColabFold-DiffDock model predicting the Crystal-Crystal dataset, while the highest *R*_*p*_ is found in the ColabFold-DiffDock model predicting the Crystal-Crystal dataset. Furthermore, the model trained on the ColabFold-DiffDock dataset generally performs better across three test sets than the other two scenarios, as shown in Fig. 3. Statistical tests, as illustrated in Table. S4, indicates that there are significant differences in performance metrics, RMSE, and *R*_*p*_ when comparing the Crystal-Crystal and ColabFold-DiffDock models. These differences are observed when predicting the Crystal-DiffDock and ColabFold-DiffDock test sets.

**Figure 3:**
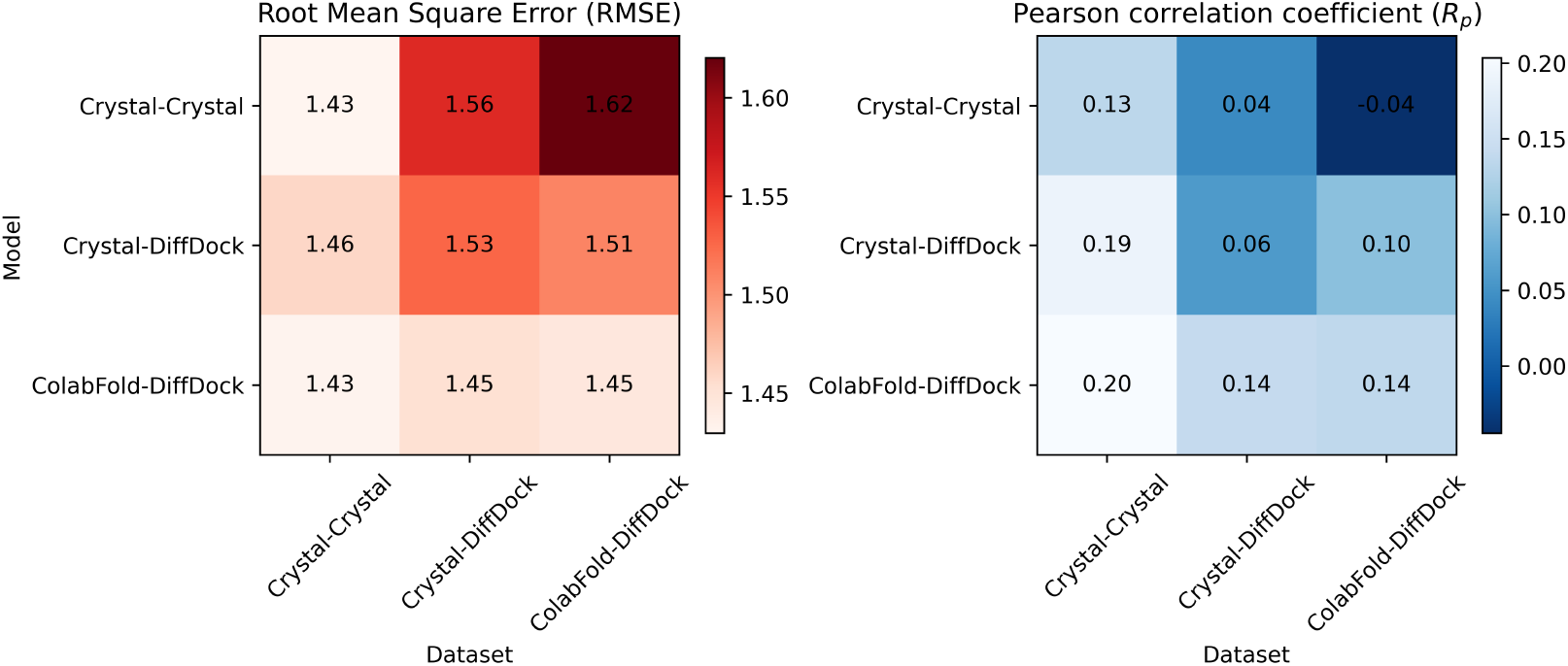
Ablation Study of Affinity Prediction Models Across Different Test Datasets. The heatmap displays the average of Root Mean Square Error (RMSE) and the average of Pearson correlation coefficient (*R*_*p*_) for Crystal-Crystal, Crystal-DiffDock, and ColabFold-DiffDock prediction models across Crystal-Crystal, Crystal-DiffDock, and ColabFold-DiffDock test datasets. We trained three distinct models and tested each of them on three separate test sets, repeating this process ten times. *R*_*p*_ and RMSE were calculated based on the predicted and true *pK*_*d*_ values.

Table. 1 presents the values corresponding to the diagonal line of Fig. 3. In other words, the table exclusively displays the performance of the three distinct prediction models evaluated on their corresponding datasets. Comparing the performance of ColabFold-DiffDock models with other SOTA docking-free methods, our framework generally achieves better on *R*_*p*_ and RMSE.

**Table 1:**
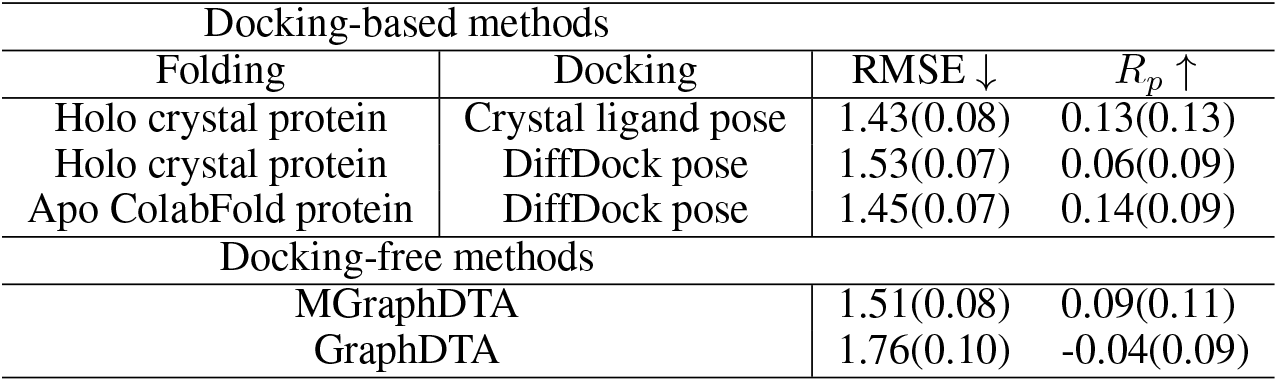
Predictive Performance of Docking-Based and Docking-Free Models in Ablation Study. The average (standard deviation) of Root Mean Square Error (RMSE) and average (standard deviation) Pearson correlation coefficient (*R*_*p*_) represent the predictive performance of three distinct FoldingDocking-Affinity (FDA) prediction models evaluated on their respective datasets. *R*_*p*_ and RMSE were calculated based on the predicted and true *pK*_*d*_ values.

| Docking-based methods |  |  |  |
| --- | --- | --- | --- |
| Folding | Docking | RMSE ↓ | $R_p$ ↑ |
| Holo crystal protein | Crystal ligand pose | 1.43(0.08) | 0.13(0.13) |
| Holo crystal protein | DiffDock pose | 1.53(0.07) | 0.06(0.09) |
| Apo ColabFold protein | DiffDock pose | 1.45(0.07) | 0.14(0.09) |
| Docking-free methods |  |  |  |
| MGraphDTA |  | 1.51(0.08) | 0.09(0.11) |
| GraphDTA |  | 1.76(0.10) | -0.04(0.09) |

In our ablation study, we tested three distinct scenarios within our FDA framework. To our surprise, the ColabFold-DiffDock combination model demonstrated a superior performance across three distinct test sets. To better investigate these counter-intuitive results, we sought to quantify the deviation introduced by ColabFold and DiffDock. We calculated the structural RMSE of the protein binding pocket structure and the ligand pose. The results of these calculations are presented in Table. 2. In the Crystal-Crystal combination, both the RMSE of the protein binding pocket structure and ligand poses are equal to zero. In the Crystal-DiffDock combination, only DiffDock is used to find the binding pocket and ligand binding pose, while the protein structures remain in their crystallized coordinates. Therefore, only the RMSE of the ligand binding poses is calculated. In the ColabFold-DiffDock combination, where both ColabFold and DiffDock are simultaneously utilized to generate binding conformations, both the RMSE of the protein binding pocket structures and ligand binding poses are calculated. When comparing the Crystal-DiffDock and ColabFold-DiffDock combinations, we observed that using DiffDock to dock ligands onto ColabFold-generated structures resulted in a higher average RMSE and standard deviation of ligand binding poses compared to using crystallized protein structures. This trend aligns with the findings from DiffDock’s work, where they tested their method on ESMfold-generated proteins (36).

**Table 2:**
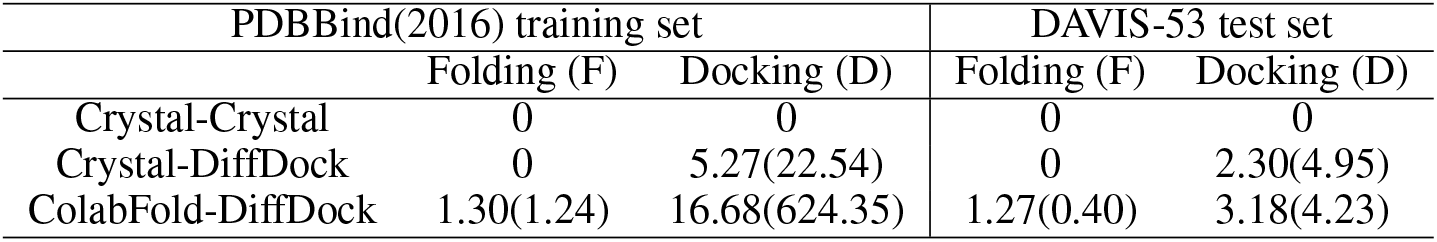
Structural RMSE Across Folding and Docking Stages. The table reports the average (standard deviation) structural Root Mean Square Error (RMSE) in the unit of Å for protein binding pocket and ligand binding poses introduced by ColabFold and DiffDock in the three distinct scenarios. Structural RMSE was calculated by comparing the predicted structures with co-crystallized experimentally determined ones.

The structural deviation analysis aligns with our hypothesis that incorporating AI models into our framework would introduce additional noise to the binding structure. However, it still does not shed some light on the observed counter-intuitive results. Further exploration of potential explanations will be presented in Discussion 3.2.

### 2.4 Binding poses augmentation

During the folding process, ColabFold has generated multiple protein conformations for each protein, and DiffDock has provided various protein-ligand binding poses for each pair during the docking process. In the original framework, the top-ranked protein conformation is selected for the docking, and the top-ranked binding pose is subsequently chosen for binding affinity prediction. This raises an interesting question: Can the lower-ranked protein conformation and binding poses be leveraged as a data augmentation strategy to enhance affinity prediction performance? To answer this question, we designed an experiment to compare the performances of different data augmentation scenarios on DAVIS and KIBA dataset with distinct data splits: (1) F-D-A: The vanilla version of our pipeline, which was implemented in Results 2.2. (2) F-5D-A: The top five binding poses generated by DiffDock are used to augment the training set. (3) F-10D-A: The top ten binding poses generated by DiffDock are used to augment the training set. (4) 2F-5D-A: For each protein-ligand pair, five binding poses were selected from both the rank-1 and rank-2 protein conformations generated by ColabFold, resulting in a total of 10 distinct binding poses. (5) 3F-5D-A: For each protein-ligand pair, five binding poses were selected from the rank-1, rank-2, and rank-3 protein conformations generated by ColabFold, resulting in a total of 15 distinct binding poses. For the test set, binding pose augmentation was not applied.

The results of the experiments are presented in Fig. 4 with KDBNet performance shown as a reference. The detailed numerical data can be found in Table. S5-S6. A comparison between the F-D-A and F-5D-A scenarios shows that augmenting the training data sets with the top five binding poses generated by DiffDock leads to an average performance increase of 12.86% in *R*_*p*_ and 8.10% in MSE on the DAVIS dataset, as well as 6.41% in *R*_*p*_ and 4.69% in MSE on the KIBA dataset. To further assess the effect of binding pose augmentation could extend, we evaluated three additional scenarios: F-10D-A, 2F-5D-A, and 3F-5D-A. In the DAVIS dataset, under the new-drug split, predictive performance improves as the number of augmented binding poses increases, with the 3F-5D-A scenario yielding a 26.71% increase in *R*_*p*_ and an 11.43% improvement in MSE. On the other hand, the new-protein and seq-id split exhibit performance saturation, suggesting limited benefits from additional binding poses. Interestingly, in the both-new split, F-5D-A scenario achieves the best performance, while performance declines as the number of augmented binding poses increases, with the 3F-5D-A scenario performing worse than the baseline (F-D-A). In the KIBA dataset, plateau effects were observed in all the data-split, except for the MSE metric in the seq-id split, which did not show improvement. These results suggest that pose augmentation enhances predictive accuracy; however, the benefits of data augmentation gradually diminish as the number of binding poses increases in most cases.

**Figure 4:**
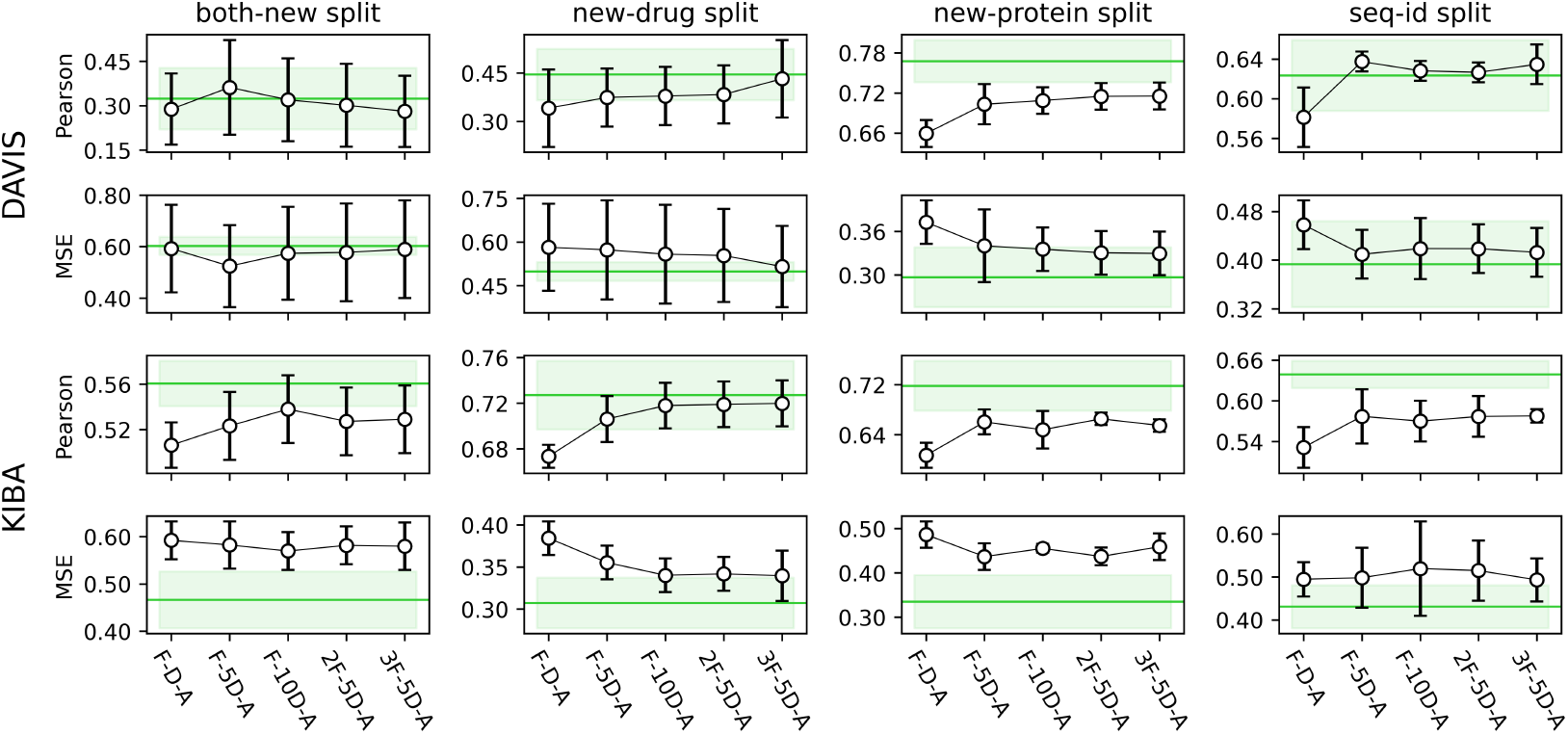
Comparison of Binding Pose Augmentation Scenarios on DAVIS and KIBA Datasets. Dot plots display the average *±* standard deviation of the evaluation results across five randomly chosen train/test splits. The light green horizontal line and area represents the average *±* standard deviation of KDBNet. Pearson correlation coefficient (*R*_*p*_) and Mean Squared Error (MSE) were calculated based on the predicted and true *pK*_*d*_ values. The detailed numerical data can be found in Table. S5-S6.

## 3 Discussion

This study mainly answers the question of whether a docking-based model surpasses a docking-free model in predicting protein-ligand binding affinity when high-resolution binding structures are unavailable. Based on our experiment result, our method generally exhibit performance comparable to the SOTA docking-free models. Additionally, it shows approximately a 32% improvement in the *R*_*p*_ for the DAVIS dataset and a 12% improvement for the KIBA dataset in the challenging ‘both-new’ split, compared to the SOTA docking-free model, MGraphDTA. These results highlight the potential generalizability of our approach to novel protein-ligand pairs. However, the advantage of the computed binding structure is not significant when accounting for the error bars. Also, there are scenarios where MGraphDTA outperforms our method in the new-protein and sequence-identity splits within the KIBA dataset. Therefore, although our docking-based models are showing promising results, the conclusion to this research question still remains to be determined. In this section, we will discuss the potential factors that could affect FDA performance and its limitations.

### 3.1 DAVIS dataset

DAVIS comprises 442 proteins, 72 ligands, and 31,824 binding affinity measurements (*K*_*d*_) but without high-resolution binding structures. It is worth noting that the truncated *K*_*d*_ (*K*_*d*_ values are greater than 10 μM) make up the majority of the DAVIS dataset (around 70%), which leads to challenges in the model training. Meanwhile, we have identified proteins within the dataset that exhibits modifications, such as mutations and phosphorylations. Previous studies (10; 11; 13; 9) tend to overlook the impact of these modifications, treating modified and unmodified proteins as equivalent. We argue that such modifications could significantly influence binding affinity predictions, and including them indiscriminately is not a prudent approach to leverage the dataset. At present, our FDA framework cannot handle such modifications. To address this limitation, we plan to utilize the AlphaFold 3 model (42) to generate the initial three-dimensional protein structure with the modification. Subsequently, we will apply biomolecular force fields to further optimize the structure and employ molecular dynamics techniques to identify its stable conformation. We anticipate that this approach will enable us to develop a more comprehensive dataset.

Secondly, given that the DAVIS dataset only provides Uniprot IDs for protein information and lacks details on the protein binding site, selecting appropriate protein structures presents a challenge. In our current work, we chose to directly utilize pre-defined PDB structures, as in the work of KDBnet (14), to represent three-dimensional structures of kinases. However, it should be noted that only half of the kinases in DAVIS have defined PDB structure, resulting in the under-utilization of the remaining data. An alternative approach involves incorporating the entire amino acid sequence of a Uniprot ID to construct the corresponding three-dimensional protein structure, which can be obtained from the Alphafold database (43). However, this method significantly expands the search space of DiffDock and increases the complexity of identifying the correct binding pockets. As a result, the predictive accuracy of ligand binding poses decreases compared to using predefined PDB structures, illustrated in Fig. S2. Therefore, we believe that constraining the protein search space is necessary in our FDA framework.

In this study, we propose a method to narrow the search space of docking by excluding unstructured protein regions. Specifically, we omit residues with model confidence scores (pLDDT) below 50 and subsequently remove residue segments shorter than 10 residues from the full-sequence AlphaFold structure. This technique yields a modest improvement over the original performance. However, it is still less effective compared to using protein amino acid sequences directly obtained from the RCSB database. The modest gains achieved with this approach suggest that while filtering out low-confidence and short segments can enhance model accuracy, the inherent quality and completeness of sequences from the RCSB database provide a more robust foundation for predictive modeling.

Despite the inherent challenges of the DAVIS dataset, it offers a valuable resource by including several protein targets with mutations and phosphorylation and their corresponding binding affinity. An interesting next step is to check if affinity predictors can discern the affinity difference between these modifications. Importantly, past studies have overlooked these changes, emphasizing the necessity of paying closer attention to the DAVIS dataset in future research. The dataset’s inclusion of various modifications provides a chance to deepen our knowledge of protein-ligand interactions and improve predictive models to handle these variations.

### 3.2 The effect of noise

In our ablation study, the model trained on apo ColabFold proteins with DiffDock-generated poses, appears to be particularly interesting. Initially anticipated as potentially the worst performing model due to the introduction of noise by ColabFold and DiffDock as shown in Table. 2, surprisingly, it generally demonstrated better performance across all three test sets. Furthermore, in our binding pose augmentation experiment, predictive performances can be enhanced by augmenting more generated binding poses into training set. We posit that through the addition of noise to binding conformations by ColabFold and DiffDock, these models seem to learn from a smoother affinity landscape toward various binding conformations. This could be advantageous for the prediction of binding affinity in various situations.

Despite observing a consistent trend in our benchmark study, ablation study, and binding pose augmentation experiments, the potential underlying mechanism remains unclear, necessitating further investigation and experimental exploration. Our analysis suggests that the noise introduced into protein structure can be separated into two distinct components: (i) The structural difference between the apo protein and the holo protein. (ii) The noise introduced by ColabFold in predicting the apo protein. Similarly, when examining the noise associated with the ligand, it can be broken down into two components: (i) Docking the ligand to apo protein instead of the holo protein. (ii) The intrinsic uncertainty of DiffDock.

In our future work, we plan to explore the structural differences between the apo protein and the holo protein. Additionally, we aim to manually introduce noise into the binding conformation by slightly perturbing the torsion angles of ligands and the torsion angles of the protein backbone and sidechains within the binding pocket area. Determining the optimal distribution of this introduced noise will be a focus of our future research.

Given that most PDB structures in the RCSB database (44) only include a subset of the complete amino acid sequence of the protein, we conducted tests using the complete-sequence AlphaFold-generated protein structures to assess performance differences. Surprisingly, we observed that this approach does not improve the performance; rather, it diminishes it, as shown in Fig. S2. Our analysis suggests that this method significantly broadens the search space of DiffDock, consequently amplifying the complexity of identifying the correct binding pockets. This leads to an introduction of excessive noise into the binding conformation.

As a result, our primary aim is to investigate how adding noise strategically can optimally enhance the prediction of binding affinity. We aim to uncover the subtle ways in which noise can influence and potentially improve the accuracy of affinity predictions, thereby shedding light on the intricate dynamics of molecular interactions within the binding pocket in the context of deep learning.

### 3.3 Limitations

While our proposed FDA method demonstrates performance comparable to SOTA docking-free models, it is essential to acknowledge its limitations. Firstly, even though our proposed method can be initiated without the necessity of three-dimensional protein and ligand structures, the generalizability of this approach across various protein types has not yet been comprehensively validated. To achieve this, our approach should be benchmarked against datasets that encompass a wide range of protein types, rather than being confined to kinase proteins alone. The challenge, however, is the lack of a comprehensive dataset that includes multiple protein types, free from overlaps with existing training data, and appropriate for effective benchmarking of our proposed method. Additionally, it also emphasizes the need for future work to develop such datasets.

Secondly, it is important to acknowledge that the training sets of ColabFold and DiffDock might contain data points that overlap with those used in our validation process. To address this, we conducted a comprehensive protein and ligand similarity analysis between these datasets. For ColabFold, which utilizes AlphaFold-Multimer, protein structures in the PDB with a release date up to April 30, 2018, were used to train the model. We performed a protein sequence identity analysis, implemented by MMseqs2 (version 13.45111) (45), between these protein structures and those from the DAVIS and KIBA datasets used in our validation process. The results, as shown in Fig. S5 and Table. S7, indicate significant protein structure overlap. Similarly, we conducted a protein sequence identity analysis between the protein structures used in the DiffDock training set and those from DAVIS and KIBA. The analysis, as shown in Fig. S6, revealed that over half of the proteins in DAVIS and KIBA are present in DiffDock’s training set. Additionally, we utilized RDKit (version 2024.03.2) to implement a ligand Tanimoto similarity analysis, which showed that a small portion of the ligands from DAVIS (8 out of 64) and KIBA (45 out of 2086) have been seen in DiffDock’s training set (Fig. S7). This overlap could potentially influence the performance metrics and lead to an overestimation of our method’s predictive capabilities. While some levels of protein overlap and ligand overlap exist, the exact protein-ligand pair overlap with ColabFold and DiffDock train sets were minimal: DAVIS has 4 out of 14,464 pairs, and KIBA has 7 out of 89,957 pairs (Table. S7). Therefore, we estimate the impact of these overlaps is probably not too substantial. However, a thorough and objective evaluation with datasets that are completely non-overlap with these training sets is difficult, because it requires carefully designed experiments or even new biochemical measurements.

Thirdly, within our FDA framework, we generate multiple protein structures during the folding process and various binding structures during the docking process. In this methodology, we followed the ranking strategy utilized in AlphaFold-Multimer (28) to rank the generated protein structures. Subsequently, the top-ranked structure was selected for the downstream docking step. During the docking process, a trained confidence model was employed to rank the generated binding poses. Consistent with the protein structure selection, the top-ranked binding pose was selected for the final binding affinity prediction. This process raises pertinent questions regarding the influence of the selection criteria on the quality of the outcomes. Specifically, how does the selection of different ranked protein structures affect the quality of the binding poses? Furthermore, how does the ranking of these binding poses impact the accuracy and reliability of the binding affinity predictions? These questions underscore the importance of the selection criteria employed at each stage of the process and its optimization. Additionally, in this study, we exclusively utilized ColabFold (31), DiffDock (36), and GIGN (25) to develop our FDA framework. The question of how other folding models, such as AlphaFold (28), ESMFold (29), and RosettaFold (34), or other docking models, such as Vina (46) and TANKBind (38), would affect the quality of binding poses and the accuracy of binding affinity predictions remains unanswered in this study. Hence, it is crucial to undertake a comprehensive investigation into these aspects to ensure the robustness and reliability of our methodology. Nevertheless, the primary objective of the present study is to evaluate the feasibility and effectiveness of our proposed framework. Thus, a thorough investigation into the selection criteria and model selection at each step remains beyond the scope of this work and will be addressed in future studies.

Lastly, our method’s explicit incorporation of the processes of protein folding and molecular docking, while enhancing predictive accuracy, incurs a substantial computational cost. The computational runtime of our method increases exponentially relative to docking-free approaches. Table. S8 and S9 presents a comparison of the average runtime and GPU memory usage for predicting binding affinity for a single protein-ligand complex between our docking-based method and docking-free ones. In our experimental evaluation, the average runtimes for the protein folding and molecular docking processes were approximately 540 seconds and 12 seconds, respectively. The affinity prediction step took an average runtime of 0.01 seconds, comparable to that of docking-free methods. However, although the process of protein folding significantly dominates the overall computational runtime, in many instances, three-dimensional predicted protein structures are readily accessible through resources such as the AlphaFold database (43). Consequently, it is often unnecessary to perform folding for each protein-ligand pair. Furthermore, for the docking part, several methods have been proposed to speed up the inference process of diffusion models. The strategies, including optimization of Stochastic Differential Equation (SDE) solvers (47), reduction of diffusion sampling time (48; 49), and the implementation of parallel processing techniques, have potential to enhance the efficiency of our method. By integrating these acceleration techniques, we anticipate that our approach could become more practical and feasible for large-scale applications.

## 4 Conclusions

In this study, we introduce a flexible framework, Folding-Docking-Affinity (FDA), for predicting protein-ligand binding affinity. This framework leverages explicitly computed binding poses, and features with interchangeable components, ensuring its adaptability to the rapidly evolving folding, docking, and affinity prediction methods. Our experiments show that the proposed framework generally exhibit performance comparable to SOTA docking-free models. Hence, the docking-based models are potentially promising, but the question of whether directly modeling 3D binding poses is necessary for accurate binding affinity prediction remains unresolved. Recent advancements in more innovative deep learning-based models, such as the updated versions of DiffDock, DiffDock-L (50), and DynamicBind (51), are noteworthy. DiffDock-L aims to improve generalizability across different types of proteins, while DynamicBind accounts for the flexibility of protein structures, which are often treated as rigid bodies in other deep learning methods. We anticipate that these advancements in molecule docking will potentially enhance the performance of our proposed framework. Additionally, our exploration led to an unexpected discovery: the introduction of noise by ColabFold and DiffDock appears to have a beneficial impact on binding affinity prediction and generalizability across diverse datasets. This interesting finding suggests that the strategic incorporation of noise may hold promise for improving the robustness and performance of binding affinity predictors. We suggest that high-resolution binding conformations are not necessarily crucial for affinity prediction; Instead, introducing moderate noise could potentially enhance the prediction performance. Moving forward, we aim to investigate the mechanisms underlying the impact of noise on model performance, as well as explore additional strategies to further refine and scale up our FDA approach. We expect that this study, could serve as a starting point for research into incorporating binding structures to improve the accuracy of binding affinity prediction.

## 5 Methods

### 5.1 Models

We employed ColabFold (v1.5.5) (31) as our folding component to generate apo protein structures. The input comprises the MSA feature generated through MMseqs2 (version 15.6f452) (45) and template structures. The backend model architecture is Alphafold 2 multimer (v3)(28). We performed three prediction cycles for each structure and generated five structures for each protein. Regarding the selection of protein structures, we followed the same ranking strategy that utilized in AlphafoldMultimer. Specifically, we ranked the generated protein structures based on a score, calculated as interface predicted template modeling score (ipTM) weighted at 0.8, and predicted template modeling score (pTM) weighted at 0.2. This ranking strategy was consistently applied across all proteins, and no fixed threshold score was implemented. The top-ranked structure was subsequently relaxed using Amber, implemented in OpenMM (7.7.0) (52).

We utilized DiffDock (v1.0) (36), a score-based diffusion model, as the docking component. For each protein-ligand pair, we sampled ten binding poses. DiffDock comprises two parts: a diffusion generative model for ligand binding poses and a confidence score model. The former samples various binding poses by generating translational and rotational matrices, along with scalars representing the torsion angles of the ligand. Additionally, the confidence score model evaluates the quality of the binding pose sampled by the generative model. One key difference between the generative model and the confidence model is that the generative model only considers the alpha-carbon of the protein structure, whereas the confidence score model takes into account all heavy atoms of the protein. We directly used the pre-trained DiffDock models for the generation and subsequent ranking of the docking poses.

Furthermore, we incorporated the affinity predictor proposed by GIGN(Git hash: ef514d3)(25) to predict binding affinity based on the top-ranked generated binding poses. GIGN incorporates three essential components: the protein graph within the binding pocket area, the ligand graph, and the protein-ligand interaction graph. Additionally, during the graph message passing phase, the model treats covalent interactions and non-covalent interactions separately to enhance the effectiveness of learning node representations.

### 5.2 Dataset

We chose two public datasets, DAVIS (40) and KIBA (41) to evaluate the effectiveness of our framework due to widespread use in benchmarking the performance of binding affinity predictors (10; 11; 12; 13; 14). It’s important to note that while the dataset provides binding affinity measurements, it lacks high-resolution crystallized binding structures. We adopted a similar method as in the KDBNet study (14) to select representative PDB structures of kinases. Ligands and kinases that do not have 3D structures in the PubChem (53) or PDB (44) databases were removed from both datasets. For kinases with several mutations in the DAVIS dataset, we retained only those with the minimal *K*_*d*_ values. The processed version includes 226 proteins, 64 ligands, and 14,464 binding affinity measurements. Additionally, we transformed the raw *K*_*d*_ values into *pK*_*d*_ values for model training. This transformation is defined as *pK*_*d*_ = *−*log_10_(*K*_*d*_*/*10^9^), which aids in model convergence by ensuring numerical stability. The KIBA dataset developed a score that integrates three distinct bioactivity parameters: *K*_*i*_, *K*_*d*_, and IC_50_. The processed version includes 160 proteins, 2,086 ligands, and 89,957 binding affinity measurements.

We also follow the KDBNet study to establish four train-test split settings to assess prediction performance: (1) new-protein split: 20% of kinases were held back as the test set, mimicking a drug repositioning scenario where the model makes predictions for unseen proteins. (2) New-drug split: 20% of drugs were set aside as the test set, emulating a drug discovery scenario where the model makes predictions for unseen drugs. (3) Both-new split: 20% of kinases and 20% of drugs were held out as the test set, representing a scenario where the model makes predictions for both unseen proteins and drugs. (4) Sequence-identity split: Similar to the new-protein split, but with the additional criterion that no kinase in the training set shares more than 50% sequence identity with any kinase in the test set.

Regarding the three-dimensional ligand structure, we directly download the ligand structure from PubChem database (53) based on the ligand’s compound identifier (CID). For three-dimensional protein structures, we leveraged the ColabFold (31) to generate the three-dimensional protein structures from the corresponding amino acid sequences. Both three-dimensional protein and ligand structures were represented as heterogeneous geometric graphs. Our node and edge feature representations follow those used in DiffDock (36).

### 5.3 Ablation study

Due to the limited availability of crystallized binding structures for most protein-ligand pairs in the DAVIS dataset, we opted to use the PDBBind (general and refined sets)(version 2016) dataset (26) as our training and validation set. We excluded protein-ligand pairs where (i) the protein with amino acid sequence longer than 3000, due to time constraints for protein folding, (ii) DiffDock failed to dock the ligand onto the protein, and (iii) DiffDock-generated ligand could not be processed by RDKit (version 2022.9.5). Ultimately, the training set consists of 11,637 protein-ligand pairs, while the validation set comprises 975 pairs. The splitting of the PDBBind dataset is applied to the docking-free models training as well.

Additionally, we identified 102 protein-ligand pairs from the DAVIS dataset that have available three-dimensional crystallized PDB structures in the RCSB database (44). To create our test set for the ablation study, we excluded 49 pairs that were already present in the DiffDock training set, resulting in a final set of 53 complex pairs (DAVIS-53) (56 unique protein chains and 16 unique ligands). Furthermore, the protein/ligand similarity analysis between PDBBind training set and DAVIS-53 was conducted by MMseqs2 (version 13.45111) (45) and RDKit (version 2024.03.2), respectively. The results of similarity distributions are illustrated in Fig. S3 and S4. Approximately 60% of the protein chains exhibit sequence identity exceeding 0.8, while 25% of the ligands display Tanimoto similarity values above 0.8.

The protein generated by ColabFold relies on the amino acid sequence extracted from the FASTA file within the RCSB database (44). This sequence serves as ColabFold’s input. Additionally, for the three-dimensional ligand structure, we employ the Tripos molecule structure format (MOL2), which is provided by PDBbind (26).

The initial step in calculating the structural RMSE for the binding pocket involves aligning the apo ColabFold protein structure with the holo crystal protein structure. For the protein generated by ColabFold, we extracted the amino acid sequence from the FASTA file in the RCSB database (44). To reconcile discrepancies such as missing amino acids in the crystal PDB file, we employed Biopython (54) for global sequence alignment, identifying the shared subsequence between the FASTA and PDB file.

Next, we utilized alpha-carbon atoms for the structure alignment, following the methodology outlined in the DiffDock work (36). This approach utilizes the weighted Kabsch algorithm. For each alpha-carbon atom *x*, its weight is described as 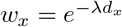 where *λ* represents a smoothing factor, and *d*_*x*_ corresponds to the minimum distance of *x* to a ligand atom within the protein-ligand complex. In this way, the alpha-carbon atom that is closer to the ligand is assigned a higher weight. Details of *λ* selection can be found in the DiffDock work (36).

After aligning the two protein structures, we calculated the RMSE using all non-hydrogen atoms within a 5 Å radius from the ligands (55), except for residues where the apo ColabFold protein and the crystallized protein have different numbers of non-hydrogen atoms.

## 6 Data availability

The processed data can be accessed at https://zenodo.org/records/15058571. This dataset includes the ColabFold-generated apo protein structures and the DiffDock-generated ligand binding poses for the affinity prediction benchmark and binding pose augmentation experiments. Additionally, for the ablation study, the repository contains three-dimensional structures of protein-ligand complexes under three scenarios (Crystal-Crystal, Crystal-DiffDock, ColabFold-DiffDock).

## 7 Code availability

The source code of the FDA is available at our GitHub repository https://github.com/ZhiGroup/FDA

## 8 Author contributions

MH.W, Z.X, and D.Z conceived the research project. Z.X, and D.Z supervised the research project. MH.W developed the computational method, implemented the software, and performed the evaluation analyses. All authors analyzed the results and participated in the interpretation. MH.W wrote the manuscript with support from all other authors.

