## Supplemental Information for "A Folding–Docking–Affinity framework for protein–ligand binding affinity prediction"

Ming-Hsiu Wu Ziqian Xie Degui Zhi
McWilliams School of Biomedical Informatics
University of Texas Health Science Center at Houston
Houston, TX, USA

#### 1 Supplementary figure

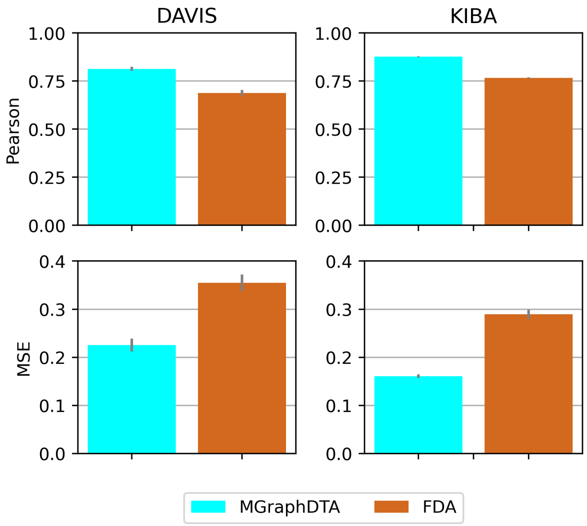


Figure S1: The performance comparison of MGraphDTA and Folding-Docking-Affinity (FDA) in the random-split of DAVIS and KIBA datasets. Bar plots display the average ± standard deviation of the evaluation results across five randomly chosen train/test splits. Pearson correlation coefficient ($R_p$) and Mean Squared Error (MSE) were calculated based on the predicted and true $pK_d$ values.

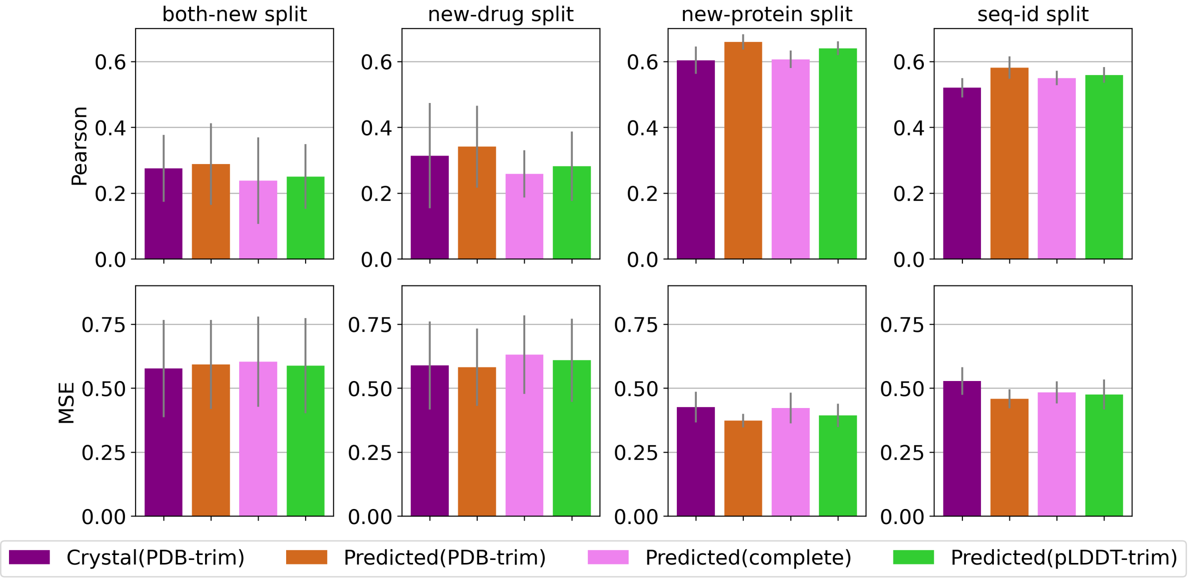


Figure S2: The performance comparison of different protein structure sources on the DAVIS dataset. Crystal(PDB-trim) refers to a protein structure obtained directly from the RCSB Protein Data Bank, which includes only a subset of the complete amino acid sequence of the protein. Predicted(PDB-trim) uses the sequence from Crystal(PDB-trim) to generate three-dimensional protein structures by ColabFold. Predicted(complete) includes the complete amino acid sequence of a UniProt ID, with the three-dimensional protein structures accessed from the AlphaFold database. Predicted(pLDDT-trim) excludes residues with model confidence scores (pLDDT) below 50 and subsequently omits residue segments shorter than 10 residues from the full Predicted(complete) structure. DiffDock and GIGN are employed for molecular docking and affinity prediction across all three scenarios. Bar plots display the average $\pm$ standard deviation of the evaluation results across five randomly chosen train/test splits. Pearson correlation coefficient ($R_p$) and Mean Squared Error (MSE) were calculated based on the predicted and true $pK_d$ values.

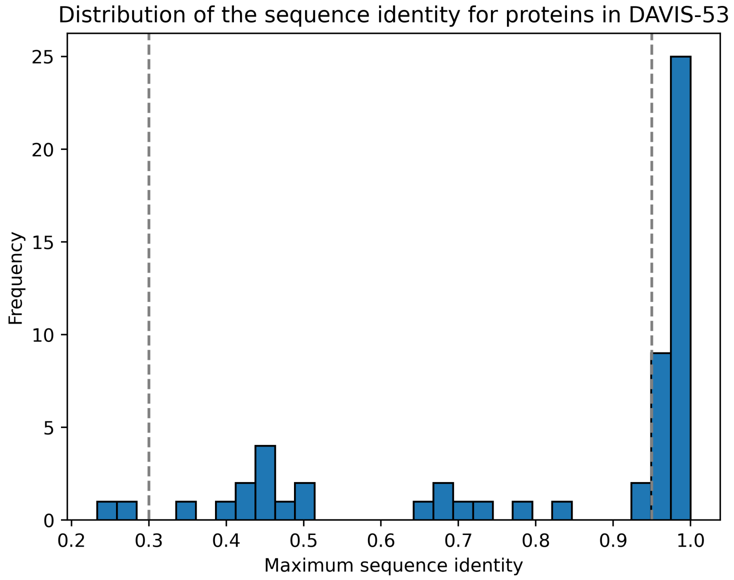


Figure S3: The distribution of protein sequence identity between DAVIS-53 and PDBBind training set. The sequence identity is the maximum sequence identity between any chains in the DAVIS-53 test protein and all PDBBind General and Refined Set v2016. The distribution of sequence identity across defined intervals is as follows: 2 protein chains fall within the range of [0%, 30%], 20 protein chains within (30%, 95%], and 34 protein chains within (95%, 100%].

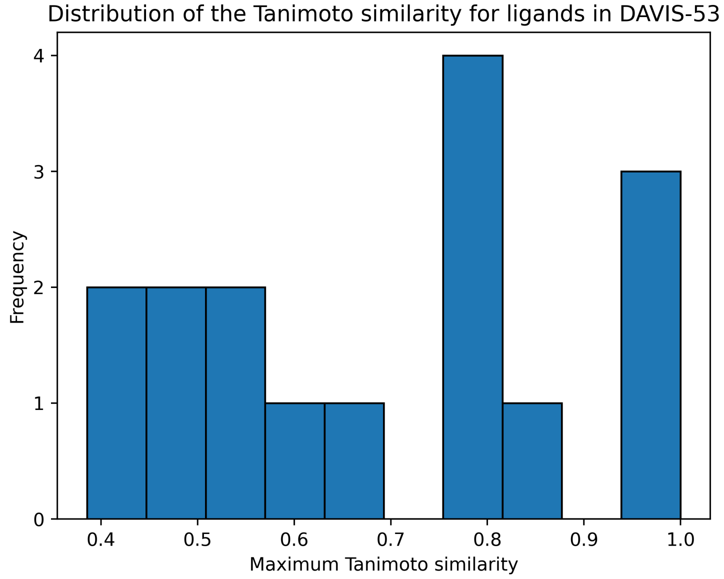


Figure S4: The distribution of ligand Tanimoto similarity between DAVIS-53 and PDBBind training set. The Tanimoto similarity is the maximum Tanimoto similarity between any ligands in the DAVIS-53 and all ligands in the PDBBind General and Refined Set v2016. In this study, molecular SMILES were converted into Morgan fingerprints, followed by the calculation of Tanimoto similarity.

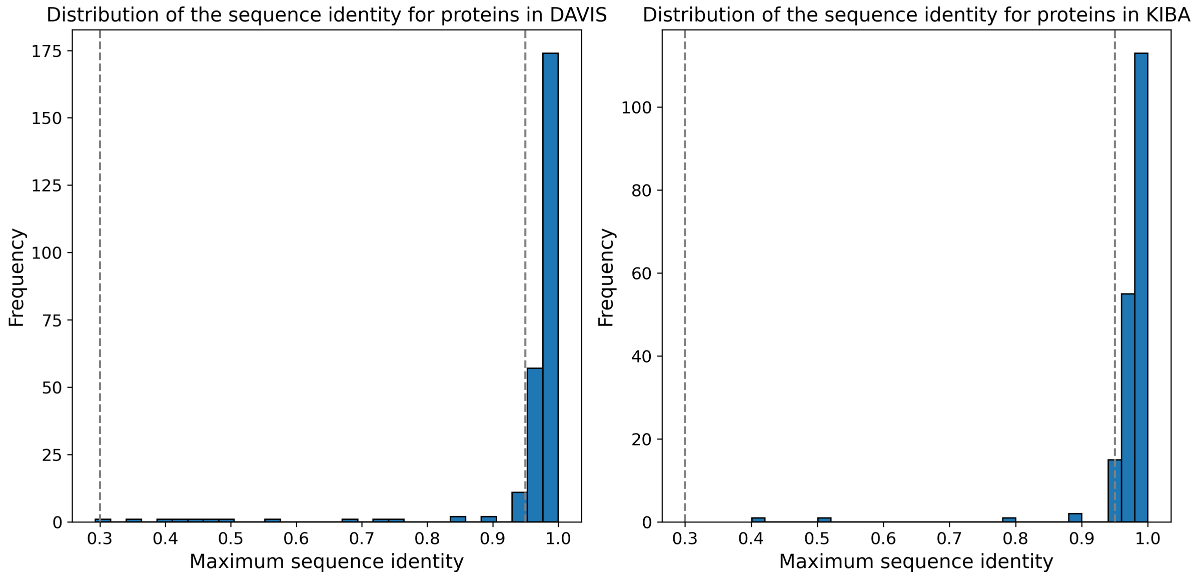


Figure S5: The distribution of protein sequence identity within the DAVIS and KIBA datasets and the AlphaFold-Multimer training dataset. Sequence identity was determined as the highest sequence identity between any protein chains in the DAVIS and KIBA datasets and all protein chains in the Protein Data Bank (PDB) as of April 30, 2018, which were used to train AlphaFold-Multimer. The sequence identity distribution across specified intervals is detailed as follows: For the DAVIS dataset, 9 protein chains fall within the range of [0%, 30%], 23 protein chains within (30%, 95%], and 233 protein chains within (95%, 100%]. Similarly, for the KIBA dataset, 4 protein chains fall within the range of [0%, 30%], 12 protein chains within (30%, 95%], and 176 protein chains within (95%, 100%].

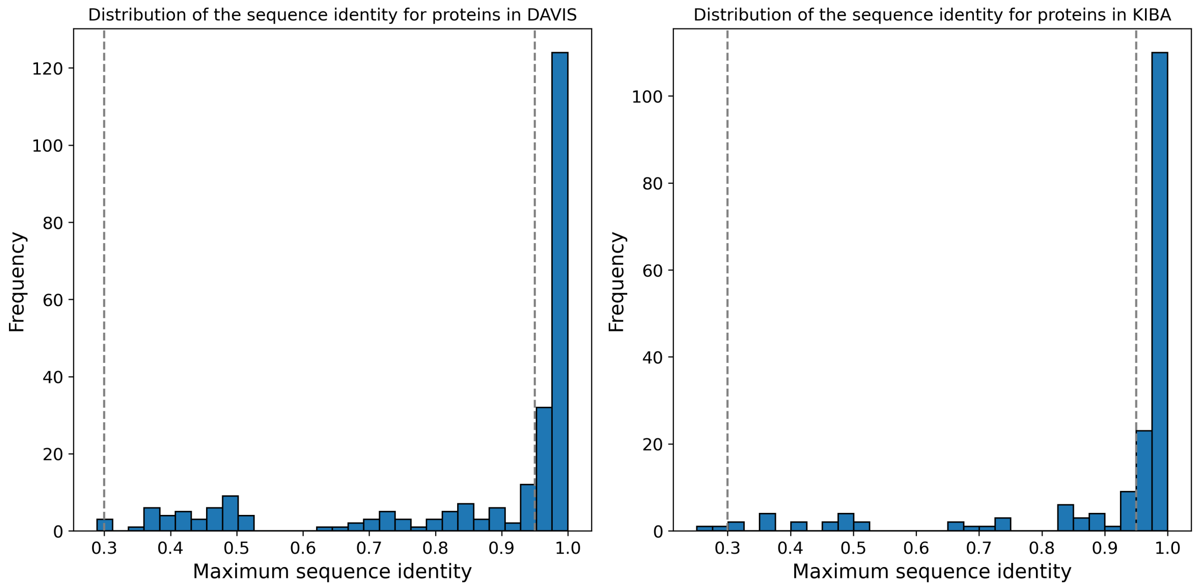


Figure S6: The distribution of protein sequence identity within the DAVIS and KIBA datasets and the DiffDock training dataset. Sequence identity was determined as the highest sequence identity between any protein chains in the DAVIS and KIBA datasets and all protein chains in the DiffDock training dataset. The sequence identity distribution across specified intervals is detailed as follows: For the DAVIS dataset, 17 protein chains fall within the range of [0%, 30%], 92 protein chains within (30%, 95%], and 156 protein chains within (95%, 100%]. Similarly, for the KIBA dataset, 13 protein chains fall within the range of [0%, 30%], 46 protein chains within (30%, 95%], and 133 protein chains within (95%, 100%].

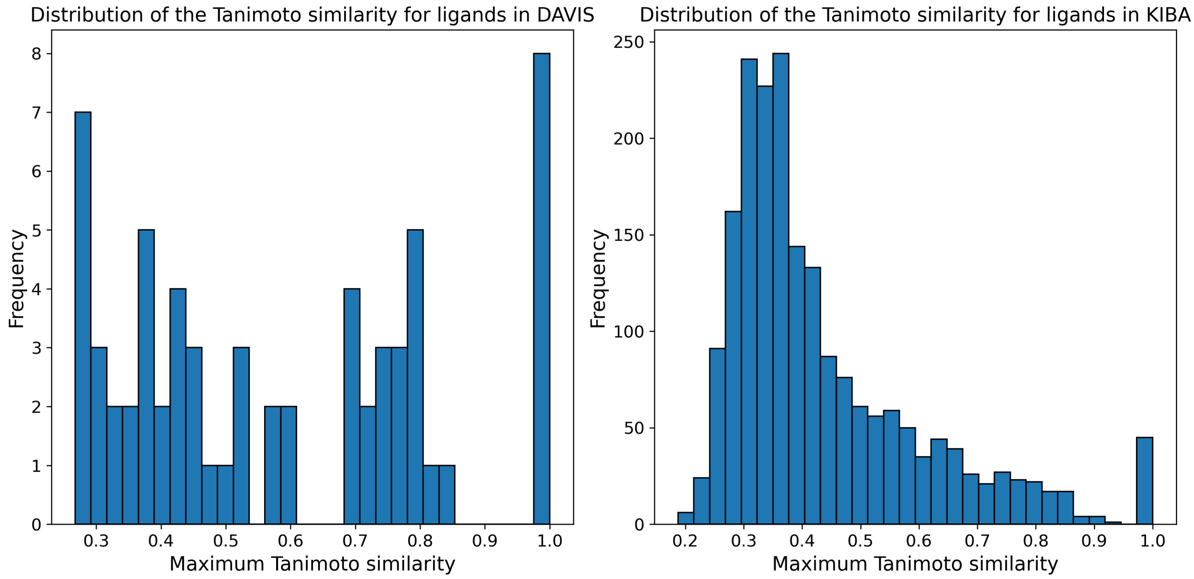


Figure S7: Distribution of ligand Tanimoto similarity within the DAVIS and KIBA datasets and the DiffDock training dataset. The Tanimoto similarity is the maximum Tanimoto similarity between any ligands in the DAVIS and KIBA datasets and all ligands in the DiffDock training dataset. In this study, molecular SMILES were converted into Morgan fingerprints, followed by the calculation of Tanimoto similarity.

### 2 Supplementary table

Table S1: Comparison of Folding-Docking-Affinity (FDA) affinity prediction performance with docking-free methods KronRLS, DeepDTA, GraphDTA, DGraphDTA, MGraphDTA, and KDBNet in four train-test split settings of the DAVIS and KIBA datasets. The average (standard deviation) of Pearson correlation coefficient ($R_p$) was calculated across five randomly chosen train/test splits. The highest $R_p$ values are shown in bold.

| | DAVIS | | | | KIBA | | | |
|---|---|---|---|---|---|---|---|---|
| | both-new | new-drug | new-protein | seq-id | both-new | new-drug | new-protein | seq-id |
| KronRLS | 0.0670(0.03) | 0.2380(0.07) | 0.4163(0.05) | 0.3611(0.08) | 0.2843(0.04) | 0.5900(0.02) | 0.5054(0.03) | 0.2775(0.10) |
| DeepDTA | 0.1238(0.04) | 0.1978(0.05) | 0.5176(0.02) | 0.4802(0.03) | 0.2858(0.05) | 0.4430(0.05) | 0.4830(0.05) | 0.3308(0.07) |
| GraphDTA | 0.0639(0.03) | 0.3471(0.07) | 0.5119(0.02) | 0.4812(0.03) | 0.3870(0.09) | 0.6188(0.07) | 0.5296(0.04) | 0.5079(0.04) |
| DGraphDTA | 0.1552(0.08) | 0.2549(0.07) | 0.5188(0.05) | 0.4996(0.03) | 0.4164(0.07) | 0.6508(0.04) | 0.5970(0.05) | 0.5598(0.03) |
| MGraphDTA | 0.2180(0.18) | 0.3381(0.11) | 0.6234(0.01) | 0.5135(0.03) | 0.4506(0.07) | 0.6476(0.02) | 0.6519(0.02) | 0.6109(0.03) |
| FDA | 0.2883(0.12) | 0.3411(0.12) | 0.6593(0.02) | 0.5813(0.03) | 0.5061(0.02) | 0.6734(0.01) | 0.6069(0.02) | 0.5312(0.03) |
| KDBNet | **0.3235**(0.10) | **0.4452**(0.08) | **0.7675**(0.03) | **0.6233**(0.04) | **0.5606**(0.02) | **0.7270**(0.03) | **0.7182**(0.04) | **0.6387**(0.02) |

Table S2: Comparison of Folding-Docking-Affinity (FDA) affinity prediction performance with docking-free methods KronRLS, DeepDTA, GraphDTA, DGraphDTA, MGraphDTA, and KDBNet in four train-test split settings of the DAVIS and KIBA datasets. Average (standard deviation) of Mean Squared Error (MSE) across five randomly chosen train/test splits. The lowest MSE values are shown in bold.

| | DAVIS | | | | KIBA | | | |
|---|---|---|---|---|---|---|---|---|
| | both-new | new-drug | new-protein | seq-id | both-new | new-drug | new-protein | seq-id |
| KronRLS | 0.9006(0.31) | 0.6852(0.15) | 0.7368(0.16) | 1.2207(0.45) | 1.1014(N/A) | 1.096(N/A) | 1.099(N/A) | 1.1050(N/A) |
| DeepDTA | 0.6344(0.08) | 0.6073(0.07) | 0.4906(0.03) | 0.4689(0.03) | 0.8036(0.10) | 0.5959(0.07) | 0.5560(0.05) | 0.7089(0.21) |
| GraphDTA | 0.8716(0.26) | 0.5455(0.06) | 0.7803(0.15) | 0.7420(0.15) | 0.6789(0.14) | 0.4831(0.08) | 0.7289(0.07) | 0.6982(0.28) |
| DGraphDTA | **0.5552**(0.06) | 0.5847(0.16) | 0.5712(0.05) | 0.4748(0.04) | 0.6125(0.05) | 0.4059(0.04) | 0.4667(0.07) | 0.5167(0.11) |
| MGraphDTA | 0.6151(0.13) | 0.5776(0.10) | 0.4064(0.02) | 0.5387(0.03) | 0.6387(0.07) | 0.4184(0.03) | 0.4396(0.03) | **0.4303**(0.04) |
| FDA | 0.5925(0.17) | 0.5816(0.15) | 0.3726(0.03) | 0.4581(0.04) | 0.5923(0.04) | 0.3842(0.02) | 0.4864(0.03) | 0.4946(0.04) |
| KDBNet | 0.6030(0.03) | **0.4983**(0.03) | **0.2968**(0.04) | **0.3930**(0.07) | **0.4663**(0.06) | **0.3072**(0.03) | **0.3349**(0.06) | 0.4305(0.05) |

Table S3: The average (standard deviation) of Root Mean Square Error (RMSE) and the average (standard deviation) of Pearson correlation coefficient ($R_p$) for Crystal-Crystal, Crystal-DiffDock, and ColabFold-DiffDock prediction models across Crystal-Crystal, Crystal-DiffDock, and ColabFold-DiffDock test datasets in the ablation study. We trained three distinct models and tested each of them on three separate test sets, repeating this process ten times. The best-performing values are highlighted in bold.

| RMSE | | | |
|---|---|---|---|
| Model | Dataset | | |
| | Crystal-Crystal | Crystal-DiffDock | ColabFold-DiffDock |
| Crystal-Crystal | **1.43**(0.08) | 1.56(0.05) | 1.62(0.08) |
| Crystal-DiffDock | 1.46(0.06) | 1.53(0.07) | 1.51(0.05) |
| ColabFold-DiffDock | **1.43**(0.11) | **1.45**(0.08) | **1.45**(0.07) |
| $R_p$ | | | |
| Model | Dataset | | |
| | Crystal-Crystal | Crystal-DiffDock | ColabFold-DiffDock |
| Crystal-Crystal | 0.13(0.13) | 0.04(0.07) | -0.04(0.06) |
| Crystal-DiffDock | 0.19(0.08) | 0.06(0.09) | 0.10(0.07) |
| ColabFold-DiffDock | **0.20**(0.12) | **0.14**(0.11) | **0.14**(0.09) |

Table S4: P-values for Root Mean Square Error (RMSE) and Pearson correlation coefficient ($R_p$) across different models and datasets comparisons. The p-values are obtained from implementing independent samples t-tests, comparing the performance metrics RMSE and $R_p$ across different model and dataset combinations. Significant p-values (when P-value $< 0.05$) are presented in bold.

| P-value for RMSE | | | |
|---|---|---|---|
| Model | Dataset | | |
| | Crystal-Crystal | Crystal-DiffDock | ColabFold-DiffDock |
| Crystal-DiffDock vs ColabFold-DiffDock | 0.5107 | 0.0609 | **0.0309** |
| Crystal-Crystal vs ColabFold-DiffDock | 0.9739 | **0.0042** | **0.0001** |
| Crystal-Crystal vs Crystal-DiffDock | 0.4025 | 0.2850 | **0.0024** |
| P-value for $R_p$ | | | |
| Model | Dataset | | |
| | Crystal-Crystal | Crystal-DiffDock | ColabFold-DiffDock |
| Crystal-DiffDock vs ColabFold-DiffDock | 0.7719 | 0.0908 | 0.3483 |
| Crystal-Crystal vs ColabFold-DiffDock | 0.2529 | **0.0337** | **0.0001** |
| Crystal-Crystal vs Crystal-DiffDock | 0.2913 | 0.6681 | **0.0001** |

Table S5: Average (standard deviation) of Pearson correlation coefficient ($R_p$) for five binding pose augmentation scenarios—F-D-A, F-5D-A, F-10D-A, 2F-5D-A and 3F-5D-A—and for the reference model (KDBNet) tested on DAVIS and KIBA dataset with distinct data splits in binding poses augmentation studies. The average (standard deviation) of $R_p$ was calculated across five randomly chosen train/test splits. The highest $R_p$ values are shown in bold.

| | DAVIS | | | | KIBA | | | |
|---|---|---|---|---|---|---|---|---|
| | both-new | new-drug | new-protein | seq-id | both-new | new-drug | new-protein | seq-id |
| F-D-A | 0.2883(0.12) | 0.3411(0.12) | 0.6593(0.02) | 0.5813(0.03) | 0.5061(0.02) | 0.6734(0.01) | 0.6069(0.02) | 0.5312(0.03) |
| F-5D-A | **0.3616**(0.16) | 0.3743(0.09) | 0.7030(0.03) | **0.6374**(0.01) | 0.5231(0.03) | 0.7061(0.02) | 0.6603(0.02) | 0.5770(0.04) |
| F-10D-A | 0.3196(0.14) | 0.3787(0.09) | 0.7086(0.02) | 0.6281(0.01) | 0.5380(0.03) | 0.7179(0.02) | 0.6479(0.03) | 0.5699(0.03) |
| 2F-5D-A | 0.3011(0.14) | 0.3835(0.09) | 0.7148(0.02) | 0.6266(0.01) | 0.5271(0.03) | 0.7189(0.02) | 0.6653(0.01) | 0.5768(0.03) |
| 3F-5D-A | 0.2808(0.12) | 0.4322(0.12) | 0.7153(0.02) | 0.6346(0.02) | 0.5290(0.03) | 0.7198(0.02) | 0.6547(0.01) | 0.5777(0.01) |
| KDBNet | 0.3235(0.10) | **0.4452**(0.08) | **0.7675**(0.03) | 0.6233(0.04) | **0.5606**(0.02) | **0.7270**(0.03) | **0.7182**(0.04) | **0.6387**(0.02) |

Table S6: Average (standard deviation) of Mean Square Error (MSE) for five binding pose augmentation scenarios—F-D-A, F-5D-A, F-10D-A, 2F-5D-A and 3F-5D-A—and for the reference model (KDBNet) tested on DAVIS and KIBA dataset with distinct data splits in Section 3.3 (Binding pose augmentation). The average (standard deviation) of MSE was calculated across five randomly chosen train/test splits. The lowest MSE values are shown in bold.

| | DAVIS | | | | KIBA | | | |
|---|---|---|---|---|---|---|---|---|
| | both-new | new-drug | new-protein | seq-id | both-new | new-drug | new-protein | seq-id |
| F-D-A | 0.5925(0.17) | 0.5816(0.15) | 0.3727(0.03) | 0.4581(0.04) | 0.5923(0.04) | 0.3842(0.02) | 0.4864(0.03) | 0.4946(0.04) |
| F-5D-A | **0.5240**(0.16) | 0.5728(0.17) | 0.3402(0.05) | 0.4096(0.04) | 0.5824(0.05) | 0.3552(0.02) | 0.4367(0.03) | 0.4980(0.07) |
| F-10D-A | 0.5740(0.18) | 0.5580(0.17) | 0.3355(0.03) | 0.4189(0.05) | 0.5696(0.04) | 0.3401(0.02) | 0.4552(0.01) | 0.5195(0.11) |
| 2F-5D-A | 0.5774(0.19) | 0.5534(0.16) | 0.3306(0.03) | 0.4187(0.04) | 0.5816(0.04) | 0.3418(0.02) | 0.4370(0.02) | 0.5148(0.07) |
| 3F-5D-A | 0.5895(0.19) | 0.5151(0.14) | 0.3297(0.03) | 0.4127(0.04) | 0.5798(0.05) | 0.3395(0.03) | 0.4590(0.03) | 0.4930(0.05) |
| KDBNet | 0.6030(0.03) | **0.4983**(0.03) | **0.2968**(0.04) | **0.3930**(0.07) | **0.4663**(0.06) | **0.3072**(0.03) | **0.3349**(0.06) | **0.4305**(0.05) |

Table S7: The number of overlapping data points between the DAVIS and KIBA datasets vs. the ColabFold (AlphaFold-Multimer) and DiffDock training sets. An overlapping protein is defined by a sequence identity greater than 95%, an overlapping ligand is defined by a Tanimoto similarity of 1, and an overlapping protein-ligand pair is defined as having the same protein amino acid sequence and ligand SMILES. The data is presented as (the number of overlapping data points) / (the total number of data points).

| | DAVIS | | KIBA | |
|---|---|---|---|---|
| | ColabFold-training | DiffDock-training | ColabFold-training | DiffDock-training |
| protein | 233/265 | 156/265 | 176/192 | 133/192 |
| ligand | - | 8/64 | - | 45/2,086 |
| protein-ligand | - | 4/14,464 | - | 7/89,957 |

Table S8: A comparison of the average runtime for predicting binding affinity for a single protein-ligand complex was conducted between our docking-based method and other models, including Folding-Docking-Affinity (FDA), MGraphDTA, and GraphDTA. The average runtime for the protein folding process was estimated by folding 226 proteins from the DAVIS dataset, which have an average sequence length of 362 amino acids, resulting in a calculated folding time of 537.48 seconds per protein (121,471 seconds / 226 proteins). Similarly, for the estimation of the average runtime for molecular docking, we predicted 14,464 protein-ligand binding poses from the DAVIS dataset, yielding an average docking time of 11.66 seconds per pose (68,622 seconds / 14,464 poses). The Folding part was tested with Python 3.10.13 and CUDA 12.3 on Ubuntu 20.04, with access to Nvidia Tesla V100 (32GB RAM), Intel(R) Xeon(R) Platinum 8168 CPU @ 2.70GHz, and 1.5TB RAM. The Docking and Affinity parts were tested with Python 3.9.18 and CUDA 11.5 on CentOS Linux 7 (Core), with access to Nvidia A100 (80GB RAM), AMD EPYC 7352 24-Core Processor, and 1TB RAM.

| Method | Runtime (s) | | | |
|---|---|---|---|---|
| | Folding | Docking | Affinity | Total (exclude Folding) |
| FDA | 537.48 | 11.66 | 0.01 | 11.67 |
| MGraphDTA | - | - | 0.02 | 0.02 |
| GraphDTA | - | - | 0.01 | 0.01 |

Table S9: A comparison of the average GPU memory usage for predicting binding affinity for a single protein-ligand complex was conducted between our docking-based method and other models, including Folding-Docking-Affinity (FDA), MGraphDTA, and GraphDTA. The average GPU memory usage is measured by batch size equal to 1.

| Method | GPU Memory (MB) | | | |
|---|---|---|---|---|
| | Folding | Docking | Affinity | Maximum |
| FDA | 422 | 2727.88 | 23.89 | 2727.88 |
| MGraphDTA | - | - | 13.05 | 13.05 |
| GraphDTA | - | - | 3.15 | 3.15 |
